# Geographical and environmental contributions to genomic divergence in mangrove forests

**DOI:** 10.1101/2020.01.08.889717

**Authors:** Michele Fernandes da Silva, Mariana Vargas Cruz, João de Deus Vidal Júnior, Maria Imaculada Zucchi, Gustavo Maruyama Mori, Anete Pereira de Souza

**Affiliations:** Department of Plant Biology, Institute of Biology, University of Campinas (UNICAMP), Campinas, SP, 13083-863, Brazil; Center for Molecular Biology and Genetic Engineering, University of Campinas (UNICAMP), Campinas, SP, 13083-875, Brazil; São Paulo Agency for Agribusiness Technology (APTA), Piracicaba, SP, 13400-790, Brazil; Institute of Biosciences, São Paulo State University (UNESP), São Vicente, SP, 11330-900, Brazil

**Keywords:** adaptation of mangroves, coastal ecology, environmental gradient, isolation by barrier, isolation by distance, molecular ecology

## Abstract

Assessing the relative importance of geographical and environmental factors to the spatial distribution of genetic variation can provide information about the processes that maintain genetic variation in natural populations. With a globally wide but very restricted habitat distribution, mangrove trees are a useful model for studies aiming to understand the contributions of these factors. Mangroves occur along the continent–ocean interface of tropical and subtropical latitudes, regions considered inhospitable to many other types of plants. Here, we used landscape genomics approaches to investigate the relative contributions of geographical and environmental variables to the genetic variation of two black mangrove species, *Avicennia schaueriana* and *Avicennia germinans*, along the South American coast. Using single nucleotide polymorphisms, our results revealed an important role of ocean currents and geographical distance in the gene flow of *A. schaueriana* and an isolation-by-environment pattern in the organization of the genetic diversity of *A. germinans*. Additionally, for *A. germinans*, we observed significant correlations between genetic variation with evidence of selection and the influence of precipitation regimens, solar radiation and temperature patterns. These discoveries expand our knowledge about the evolution of mangrove trees and provide important information to predict future responses of coastal species to the expected global changes during this century.

## INTRODUCTION

Environmental and geographical variation often affects the distribution of allele frequencies in natural populations by facilitating or limiting gene flow across space and driving selection of certain genotypes (Wang *et al.*, 2013; Sork, 2016; Murray *et al.*, 2019). For example, under an isolation-by-distance (IBD) model (Wright, 1943), geographical distance might limit dispersal, leading to the accumulation of divergences in allele frequency by genetic drift (Bradburd *et al.*, 2013). Furthermore, under the isolation-by-barrier (IBB) model, a barrier to gene flow might abruptly reduce or even disrupt connectivity between individuals of a species (Barton, 1979). In addition to these models, genetic differentiation can increase in response to environmental differences, regardless of the geographical distance. This pattern, described by the isolation-by-environment (IBE) model (Wang & Bradburd, 2014), can be generated by a variety of ecological processes, such as selection against immigrants, leading to the evolution of locally adapted populations (Bradburd *et al.*, 2013).

The IBD, IBB and IBE models are not mutually exclusive and often co-occur in nature (Wang, 2013; Sexton *et al.*, 2014). Studies aiming to determine the factors that control the distribution patterns of genetic variation across space can provide relevant information about the underlying processes that generate and maintain genetic variation in natural populations (Lee & Mitchell-Olds, 2011; Wang & Bradburd, 2014). This knowledge is essential to predict future responses of current populations to environmental changes (Vincent *et al.*, 2013) and can contribute to decision-making processes aiming to minimize future biodiversity loss (Kovach *et al.*, 2012; Muñoz *et al.*, 2015; Wee *et al.*, 2019).

The field of research that seeks to clarify the roles of these factors in the distribution of the neutral and adaptive genetic variability of a species over space is known as landscape genomics (Joost *et al.*, 2007; Lowry, 2010; Schoville *et al.*, 2012; Vincent *et al.*, 2013). Recently, this approach has been applied increasingly to the study of non-model organisms (Storfer *et al.*, 2018). Nevertheless, landscape genomic studies are mostly limited to animal species, whereas studies on plants, especially tropical trees (Storfer *et al.*, 2010), remain very limited despite their fundamental roles in global biogeochemical cycles (Jasechko *et al.*, 2013) and as habitat providers to most terrestrial biodiversity (Mannion *et al.*, 2014).

As sessile organisms, trees respond directly to the environment in which they live (Holderegger *et al.*, 2010). Conversely, they often have high levels of gene flow, which tend to reduce the strength of natural selection (Savolainen *et al.*, 2007). Thus, tree species that often occur across wide latitudinal ranges along environmental gradients, such as mangrove forests (Tomlinson, 1986), represent a promising biological model to investigate and understand the effects of the environment on microevolutionary processes and population dynamics.

Mangrove trees occur in a narrow area along the continent–ocean interface (Tomlinson, 1986; Hamilton, 2020) of tropical and subtropical latitudes of the world, mainly between 30°N and 30°S (Giri *et al.*, 2011). These species produce floating seeds or fruits, referred to as ‘propagules’ (Tomlinson, 1986), which can disperse over long distances via rivers and ocean surface currents (Van der Stocken *et al.*, 2019b). Their geographical distribution is limited by the topography of the intertidal zone (Middleton, 2012) and, frequently, by the occurrence of low temperatures (Morrisey *et al.*, 2010) and by patterns of precipitation (Spalding *et al.*, 1997). However, environmental variations in their boundaries are associated with different climatic thresholds (Osland *et al.*, 2017; Cavanaugh *et al.*, 2018) and oceanographic conditions (Soares *et al.*, 2012; Saintilan *et al.*, 2020).

*Avicennia* L. (Acanthaceae) is one of the most diverse and widely distributed mangrove genera globally (Duke, 1991; Li *et al.*, 2016) and is highly abundant on the Western Atlantic coastline (Schaeffer-Novelli *et al.*, 1990). In this region, *Avicennia* is represented mainly by two of the three New World *Avicennia species, namely, Avicennia germinans* (L.) L. and *Avicennia schaueriana* Stapf & Leechman ex Moldenke (Schaeffer-Novelli *et al.*, 1990; Duke, 1991). These species present a partly sympatric distribution on the South American coast, where they share a remarkable north– south pattern of genetic divergence, as revealed by selectively neutral microsatellites (Mori *et al.*, 2015) and single nucleotide polymorphisms (SNPs) (Cruz *et al.*, 2019, 2020). A similar pattern of putatively neutral genetic diversity has also been observed in other coastal species, such as *Rhizophora mangle* (Pil *et al.*, 2011; Francisco *et al.*, 2018) and the mangrove- associated tree *Hibiscus pernambucensis* (Takayama *et al.*, 2008). These findings probably indicate a prominent role of the dispersal of floating propagules in shaping the overlapping north–south genetic divergence of these trees, given the bifurcation of the South Equatorial Current (SEC) along the Atlantic coast of South America (Lumpkin & Johnson, 2013).

In addition to neutral processes that shape the diversity of *Avicennia* species in this region, recent studies have identified various genomic regions that might be associated with adaptive processes relevant to the environmental context of mangroves (Cruz *et al.*, 2019, 2020). These variations have been attributed to climatic and oceanographic factors that vary widely along the latitudinal gradient of the species distribution. Although these studies provide insights into the role of the environment in the organization of the adaptive genetic variation in *A. schaueriana* and *A. germinans*, the relative importance of neutral and non-neutral abiotic factors remains unknown.

In this study, we explore the relative contributions of geographical and environmental factors to the organization of the genetic diversity of *A. schaueriana* and *A. germinans* along the Atlantic coast of South America. We collect previously published genetic information and spatial and environmental data and conduct landscape genetic analyses to assess the hypothesis that geographical distance, SEC bifurcation and climatic, oceanographic and tidal variations drive population genetic differentiation of the two species, i.e. IBD, IBB and IBE models. Considering the genetic structure inferred in previous studies (Cruz *et al.*, 2019, 2020; Mori *et al.*, 2015) and the environmental heterogeneity throughout the species distribution, we expect that IBB and IBE will be the main models underlying the geographical distance for both species. Finally, after identifying the factors influencing the distributions of the genetic diversity for *A. schaueriana* and *A. germinans*, we discuss the implications for conservation and provide suggestions to improve the long-term resilience of the two black mangrove species.

## MATERIAL AND METHODS

### Biological materials and genotyping of SNP markers

Biological materials were collected and SNP markers identified as described in previous studies by Cruz *et al.* (2019) for *A. schaueriana* and by Cruz *et al.* (2020) for *A. germinans*. Briefly, 77 *A. schaueriana* individuals were sampled from ten different locations and 48 *A. germinans* from six different locations along the Brazilian coast (Fig. 1 and Table 1). Assembly, mapping and SNP locus identification were performed using custom scripts (SNPsaurus, LLC), which created a reference catalogue of abundant reads, retaining biallelic loci present in ≥ 10% of the samples. High-quality sequences were used, allowing a maximum of 65% of missing data and one SNP per sequence and requiring a minimum coverage of 8× and a minor allele frequency ≥ 0.05 using Vcftools v.0.1.12b (Danecek *et al.*, 2011). A maximum reading cover of 56 was used (resulting from the product with the average read depth and a standard deviation of 1.5 from the average) to reduce the paralogy or low-quality genotype calls. In total, 6170 and 2297 SNP markers were identified for *A. schaueriana* and *A. germinans*, respectively.

**Figure 1.**
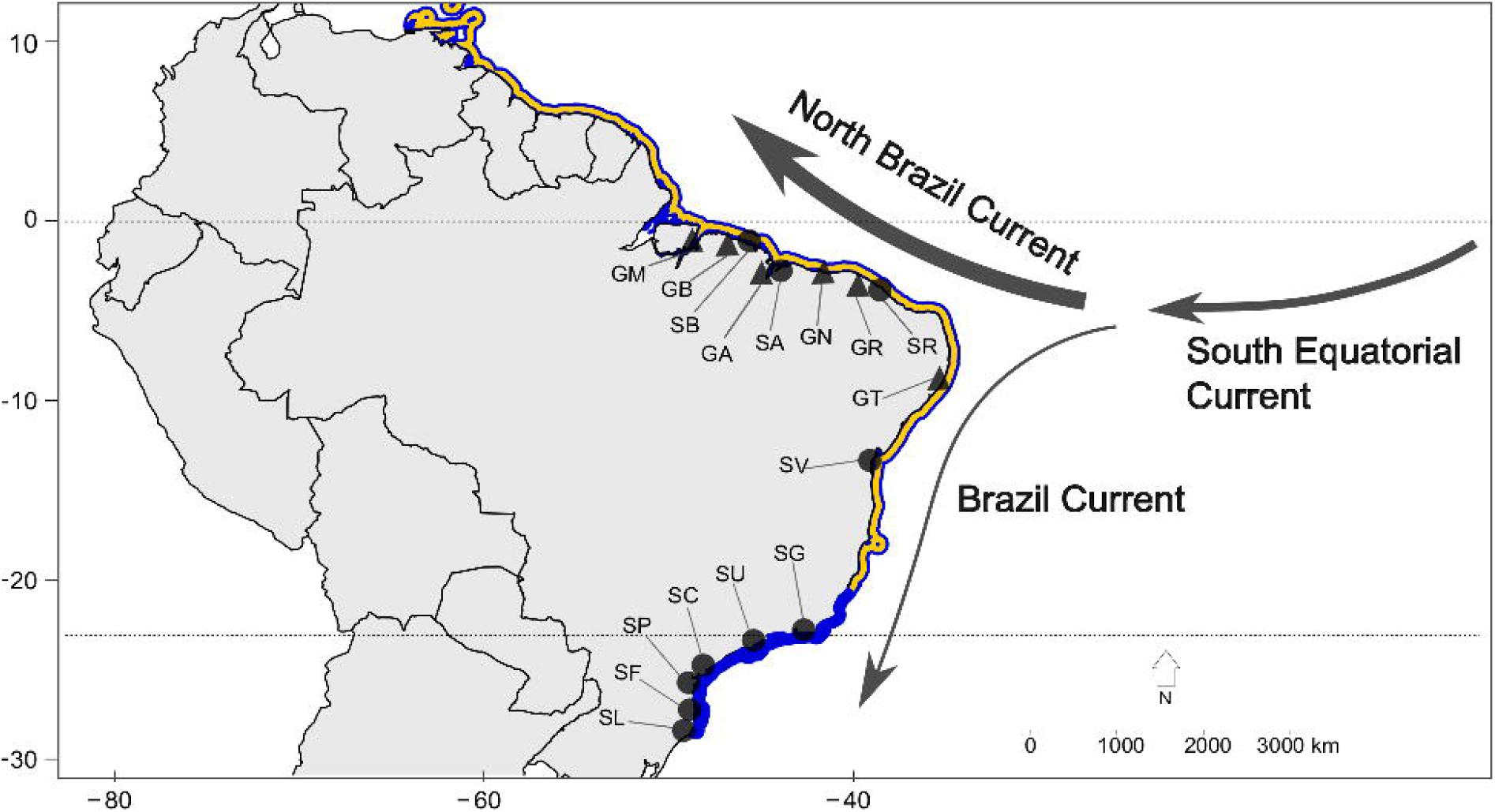
Geographical distribution and characterization of the *Avicennia schaueriana* and *Avicennia germinans* sampling sites along the South American coast. The blue area represents the geographical distribution of *A. schaueriana* and its sympatric region with *A. germinans* (blue and yellow). Black circles and triangles represent the sampling sites of the plant material used for the genotyping of genomic DNA from *A. schaueriana* and *A. germinans*, respectively (Cruz *et al.*, 2019, 2020). Sampling locations, which are indicated by two letters, are displayed as described in Table 1. Arrows represent the direction of the main ocean currents acting on the Brazilian coast. Arrow widths illustrate the mean current speed (Lumpkin & Johnson, 2013).

**Table 1.** Sites at which the samples of *Avicennia schaueriana* and *A*vicennia *germinans* were collected along the South American coast, from north to south.

| *A. schaueriana* | *A. germinans* | Locality (City/State) | Latitude | Longitude |
| --- | --- | --- | --- | --- |
|  | GM | Soure, Pará | -0.723 | -48.490 |
|  | GB | Bragança, Pará | -0.904 | -46.687 |
| SB |  | Bragança, Pará | -0.820 | -46.615 |
| SA | GA | Alcântara, Maranhão | -2.410 | -44.406 |
|  | GN | Parnaíba, Piauí | -2.778 | -41.822 |
| SR | GR | Paracuru, Ceará | -3.413 | -39.056 |
|  | GT | Tamandaré, Pernambuco | -8.526 | -35.013 |
| SV |  | Vera Cruz, Bahia | -12.983 | -38.684 |
| SG |  | Guapimirim, Rio de Janeiro | -22.701 | -43.007 |
| SU |  | Ubatuba, São Paulo | -23.489 | -45.164 |
| SC |  | Cananéia, São Paulo | -25.020 | -47.918 |
| SP |  | Pontal do Paraná, Paraná | -25.575 | -48.352 |
| SF |  | Florianópolis, Santa Catarina | -27.576 | -48.518 |
| SL |  | Laguna, Santa Catarina | -28.445 | -48.840 |

Sample populations of *A. schaueriana* and *A. germinans* are indicated by two capital letters, as shown in Figure 1.

### Detection of SNP loci with signatures of natural selection

The SNP loci with evidence of natural selection were identified previously by Cruz *et al.* (2019, 2020). For *A. schaueriana*, 86 loci showed considerable deviations from neutral expectations of interpopulation divergence. They were detected using two methods to minimize false positives: LOSITAN (Antao *et al.*, 2008), with a confidence interval of 0.99 and a false-discovery rate (FDR) of 0.1, and *pcadapt* (Luu *et al.*, 2017), with an FDR of 0.1. For *A. germinans*, 25 loci showed considerable deviations from the neutral expectations of interpopulation divergence. For the latter species, in addition to LOSITAN (a confidence interval of 0.99 and an FDR of 0.05) and *pcadapt* (an FDR of 0.05), SNP loci associated with ecological variables were detected using latent factor mixed models (LFMM) implemented in the *LEA* package (Frichot & François, 2015).

### Estimation of genetic distances

To analyse the importance of geographical distance, oceanographic barriers and environmental variables in spatial genetic divergence, we evaluated which model (IBD, IBB or IBE) best described the distribution of the genetic diversity of each species based on genome-wide SNP markers. To that end, we estimated the pairwise genetic differentiation (Wright’s *F*_ST_; Wright, 1949) for the total set of SNP molecular markers and for the set of SNP markers with evidence of selection using the *Hierfstat* package (Goudet, 2005) for R v.3.6.2 (R Core Team, 2019).

### Geographical and environmental distances and the oceanographic barrier

Pairwise geographical distances among populations were measured using the geographical coordinates of the sampling sites (Table 1) with the global positioning system (Garmin 76CSx, WGS-84 standard; Garmin International Inc., Olathe, KS, USA). Distances between points were estimated based on the contour of the Brazilian coast; thus, we considered floating propagule-mediated dispersal (Van der Stocken *et al.*, 2019a). A binary matrix (zero or one) was constructed based on the presence (one) or absence (zero) of the supposed oceanographic barrier between each pair of sampling sites to determine the relative significance of the pattern of splitting of the SEC into the Brazil Current (BC) and the North Brazil Current (NBC) (Lumpkin & Johnson, 2013) (Fig. 1) for *A. schaueriana* and *A. germinans* propagules (Cushman *et al.*, 2006; Robertson *et al.*, 2009; Legendre & Legendre, 2012; Wu *et al.*, 2016).

We obtained 42 environmental variables for each sampling site, with a resolution of 30 arc-s (~1 km in Ecuador), to evaluate the overall effect of the environment on the distribution of genetic variation. In our dataset, we included 27 climatic variables derived from the WorldClim temperature and precipitation datasets (v.1.4 and v.2.0; Fick & Hijmans, 2017), ten oceanographic variables derived the MARSPEC ocean surface salinity and temperature datasets (Sbrocco & Barber, 2013) and five variables related to tidal variations retrieved from the Environmental Climate Data Sweden (ECDS) platform (Klein *et al.*, 2013). We removed variables that showed a high correlation (r > 0.8; Supporting Information, Figure S1), as measured by the *removeCollinearity* function of the *virtualspecies* package (Leroy *et al.*, 2016) in R (R Core Team, 2019), to avoid non-independence between environmental variables. We extracted the values of the environmental variables for our sample points (Supporting Information, Tables S1 and S2) using the *raster* package (Hijmans, 2017) in R (R Core Team, 2019). For terrestrial variables, the extraction step was performed for points that overlapped our geographical coordinates. For oceanographic variables, we used a 5 km buffer around each population sampled and extracted the mean values inside the buffer; thus, non-terrestrial areas around our sampling sites were included. All occurrence data have been carefully inspected to detect and correct problems associated with inconsistent records (Chapman, 2005). We transformed this environmental data matrix using a principal components analysis (PCA). The scores for the first five and first three principal components that retained > 90% of the variance of the environmental variables for *A. schaueriana* and *A. germinans*, respectively, were used to calculate the Euclidean distances between population pairs. The PCA and environmental distance measurements were all performed in R (R Core Team, 2019).

### Association tests

We investigated the relationships between genetic differentiation (both neutral and putatively non-neutral) and geographical/environmental factors using a combination of Mantel tests (simple and partial) and matrix regression analysis. Initially, we performed Mantel tests to assess the correlations between genetic differentiation and the geographical distance, oceanographic barrier matrix and environmental distance. Next, we conducted partial Mantel tests (Smouse *et al.*, 2012) to estimate the influence of one factor conditioned to another factor as a covariate (Legendre, 1993). Both Mantel tests were conducted using the ‘*ecodist*’ package (Goslee & Urban, 2007), with 10 000 permutations.

In addition, we performed multiple matrix regression with randomization (MMRR) using the *MMRR* function in R with 10 000 permutations (Wang, 2013). We used this method to estimate the independent effect of each factor and quantify how genetic distances respond to changes in predictor variables. MMRR has proved to be accurate for several types of conditions (Wang, 2013); however, as in many multiple regression analyses, MMRR can be biased when the predictor variables are correlated (Wang, 2013). Therefore, we interpret our results based on this possible limitation.

We performed correlations for the set of all populations and also for the set of samples located northerly from the SEC and for the set of samples located southerly from the SEC because substantial variations in the genetic structure have been reported at smaller geographical scales in mangroves located in these regions (Cruz *et al.*, 2019, 2020; Mori *et al.*, 2015). Previous findings indicate that genetic diversity is organized in well-defined groups at the regional scale for both species and even between individuals of A*. germinans* that are separated by only a few kilometres (Cruz *et al.*, 2020).

Finally, given that IBE was suggested as a useful model to describe the observed genetic differentiation, we conducted an MMRR analysis and partial Mantel tests for each environmental variable separately to identify the most crucial environmental factors that affect genetic distance with evidence of natural selection.

## RESULTS

### Genetic, geographical and environmental distances

For the total set of SNP markers, we obtained *F*_ST_ values ranging from 0.019 to 0.189 for pairs of *A. schaueriana* populations and from 0.047 to 0.387 for pairs of *A. germinans* populations (Fig. 2). For the set of markers with evidence of selection, the *F*_ST_ ranged from 0.02 to 0.36 for *A. schaueriana* (Supporting Information, Table S3) and from 0.03 to 0.88 for *A. germinans* (Supporting Information, Table S4). The pairwise geographical distances ranged from ~5000 km between Bragança (SB) and Laguna (SL) to 77 km between Cananéia (SC) and Pontal do Paraná (SP) for *A. schaueriana* (Supporting Information, Table S5) and from 2100 km between Ilha de Marajó in Soure (GM) and Tamandaré (GT) to 222 km between Ilha de Marajó in Soure (GM) and Bragança (GB) for *A. germinans* (Supporting Information, Table S6).

**Figure 2.**
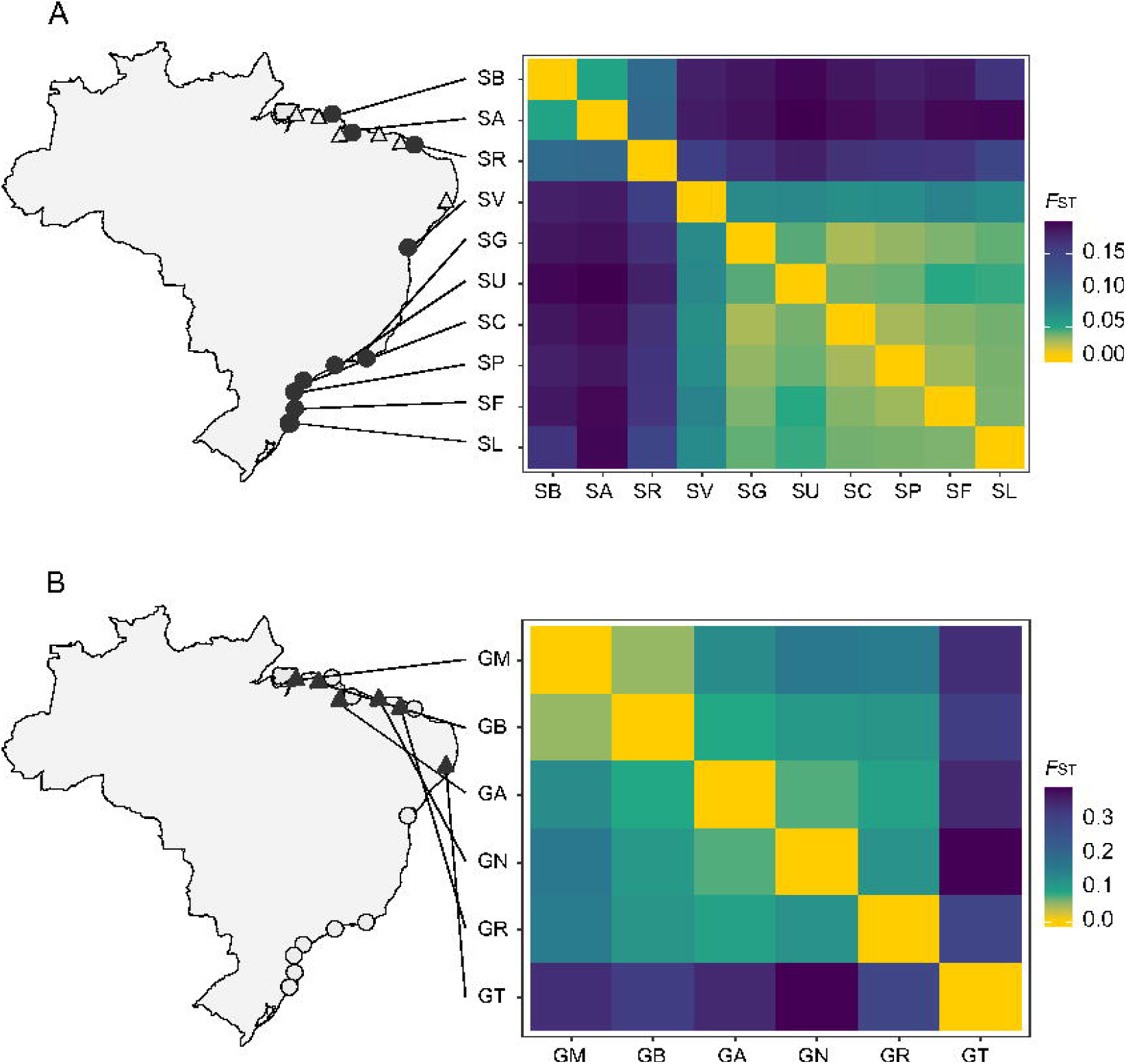
Heatmaps describing interpopulation genetic differentiation based on the total set of single nucleotide polimorphism molecular markers estimated by pairwise *F*_ST_ (Wright) for *Avicennia schaueriana* (filled circles; A) and *Avicennia germinans* (filled triangles; B) collection sites. Sample codes are denoted as in Table 1.

After removing highly correlated environmental variables (r > 0.8), 23 variables were retained for analyses of environmental distances (Supporting Information, Tables S7 and S8). The first five axes of the PCA of *A. schaueriana* retained 97% of the variance of the environmental variables used to calculate the environmental distance between the sampling sites (Supporting Information, Tables S9 and S10). For *A. germinans*, this calculation was performed based on the first three axes of the PCA, which retained 92% of the data variance. The first two PCA axes for *A. schaueriana* represented 80% of the variance and were explained mainly by the variations in air temperature and sea surface temperature, the solar radiation and tidal cycles (Fig. 3A). For *A. germinans*, the first two PCA axes represented ~80% of the variance and were explained mainly by the air temperature variation, precipitation regimens, vapour pressure deficit and solar radiation (Fig. 3B).

**Figure 3.**
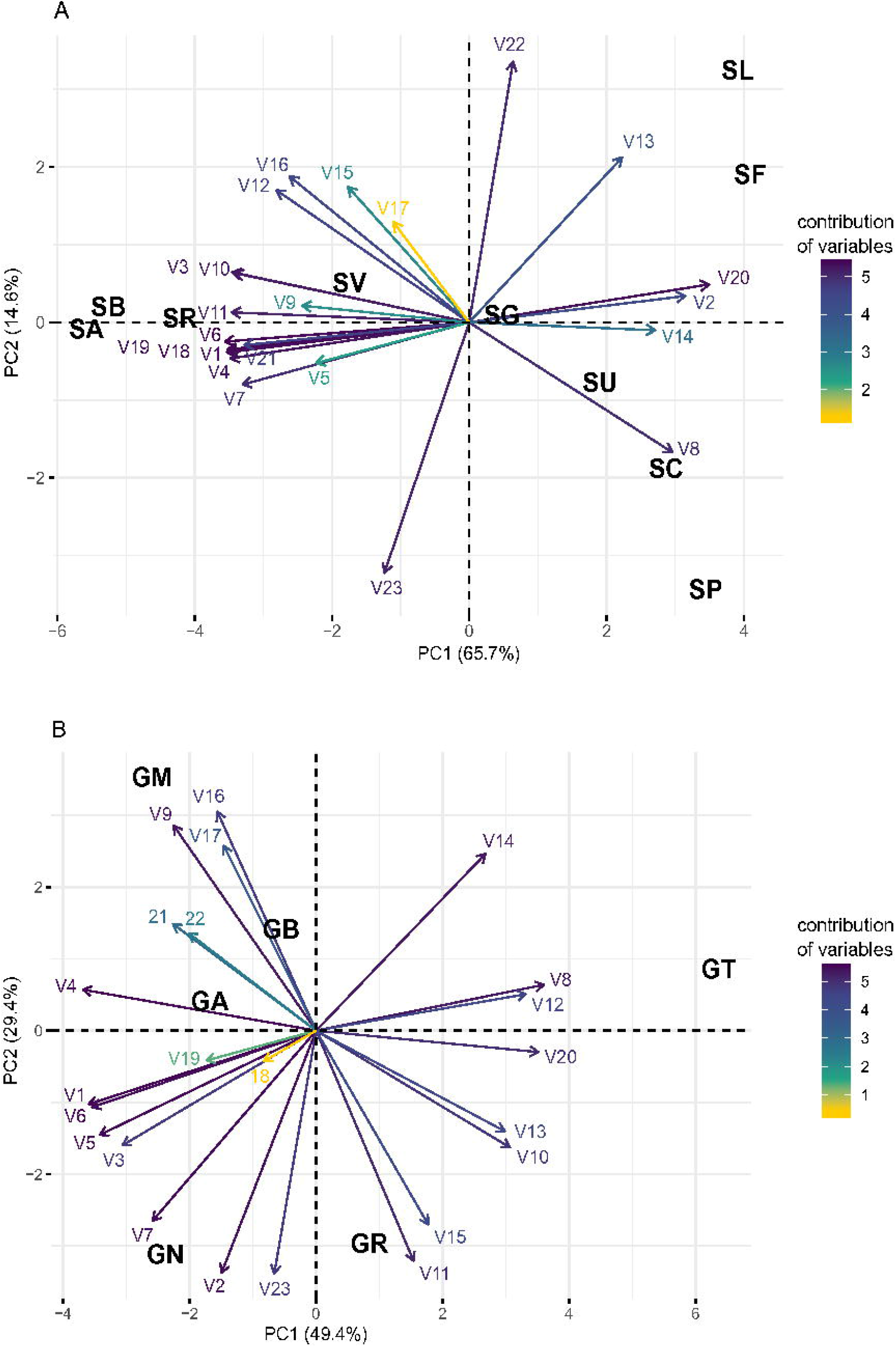
Locations of populations of *Avicennia schaueriana* (A) and *Avicennia germinans* (B) in a bidimensional projection of the principal components analisys used to calculate the environmental distance. The acronyms for sampling locations, which are indicated by two letters, are the same as those listed in Table 1. The color gradient represents the contribution of each environmental variable to principal component (PC) and PC2. Abbreviations: V1, annual mean temperature (in degrees Celsius); V2, annual temperature range (in degrees Celsius); V3, isothermality (in degrees Celsius); V4, minimum temperature of the coldest month (in degrees Celsius); V5, maximum temperature of the warmest month (in degrees Celsius); V6, mean temperature of the coldest quarter (in degrees Celsius); V7, precipitation seasonality (in millimetres); V8, precipitation in the warmest quarter (in millimetres); V9, precipitation in the coldest quarter (in millimetres); V10, mean solar radiation (in kilojoules per square metre per day); V11, minimum solar radiation (in kilojoules per square metre per day); V12, maximum solar radiation (in kilojoules per square metre per day); V13, mean wind speed (in metres per second); V14, maximum vapour pressure deficit (in kilopascals); V15, mean annual sea surface salinity (in practical salinity units); V16, sea surface salinity in the saltiest month (in practical salinity units); V17, annual variance in sea surface salinity (in practical salinity units); V18, mean annual sea surface temperature (in degrees Celsius); V19, sea surface temperature in the coldest month (in degrees Celsius); V20, annual range in sea surface temperature (in degrees Celsius); V21, anual average cycle amplitude (in centimetres); V22, annual average duration of tidal cycles (in hours); V23, annual number of cycles.

### Association tests

For *A. schaueriana*, simple Mantel tests that included all SNP loci revealed significant correlations between genetic distance and the three predictor variables, namely, geographical distance (*r* = 0.9, *P* < 0.001), environmental distance (*r* = 0.73, *P* < 0.001) and oceanographic barrier matrix (*r* = 0.96, *P* < 0.01) (Table 2 and Fig. 4). However, all predictor variables were also highly correlated with each other (geographical vs. environmental distance: *r* = 0.9, *P* < 0.001, (Fig. 4); geographical distance vs. oceanographic barrier: *r* = 0.87, *P* < 0.01; environmental distance vs. oceanographic barrier: *r* = 0.72, *P* < 0.01). When the influence of the other two factors was controlled in partial Mantel tests, the associations between genetic distance and geographical distance and between genetic distance and the oceanographic barrier matrix remained significant, whereas the correlation between genetic differentiation and environmental distance was not significant (Table 2). In addition, the multivariate regression analysis with the combined effect of the three predictor variables on the genetic distance did not show significant results for the environment (β_geographical_ = 0.48, *P* < 0.01; β_environment_ = −0.18, *P* = 0.14; β_ocean barrier_ = 0.67, *P* < 0.01; Table 3), and when this factor was removed, the oceanographic barrier variable provided a relatively higher contribution than the geographical distance (β_geographical_ = 0.25, *P* < 0.01; β_ocean barrier_ = 0.73, *P* < 0.01; Table 3). When the tests were performed separately for sampling sites located to the north and south of the SEC, we observed significant correlations only between genetic and geographical distances (partial Mantel: *r* = 0.88, *P* = 0.01; MMRR: β_geographical_ = 0.10, *P* = 0.01; β_environment_ = −0.25, *P* = 0.17) for sampling sites located south of the SEC.

**Table 2.** Results of the simple and partial Mantel tests between genetic distance based on the total set of single nucleotide polymorphism molecular markers and the geographical distance, environmental distance and oceanographic barrier matrix for *Avicennia schaueriana* populations.

| Landscape feature | Controlled | *r* | *P* |
| --- | --- | --- | --- |
| Geographical distance |  | 0.9 | < **0.001** |
| Environment distance |  | 0.73 | < **0.001** |
| Oceanographic barrier |  | 0.96 | < **0.01** |
| Geographical distance | Environment distance | 0.83 | < **0.001** |
| Geographical distance | Oceanographic barrier | 0.45 | < **0.01** |
| Environment distance | Geographical distance | - 0.47 | 0.9 |
| Environment distance | Oceanographic barrier | 0.23 | 0.09 |
| Oceanographic barrier | Geographical distance | 0.82 | < **0.01** |
| Oceanographic barrier | Environment distance | 0.92 | < **0.01** |

Significant values are presented in bold.

**Figure 4.**
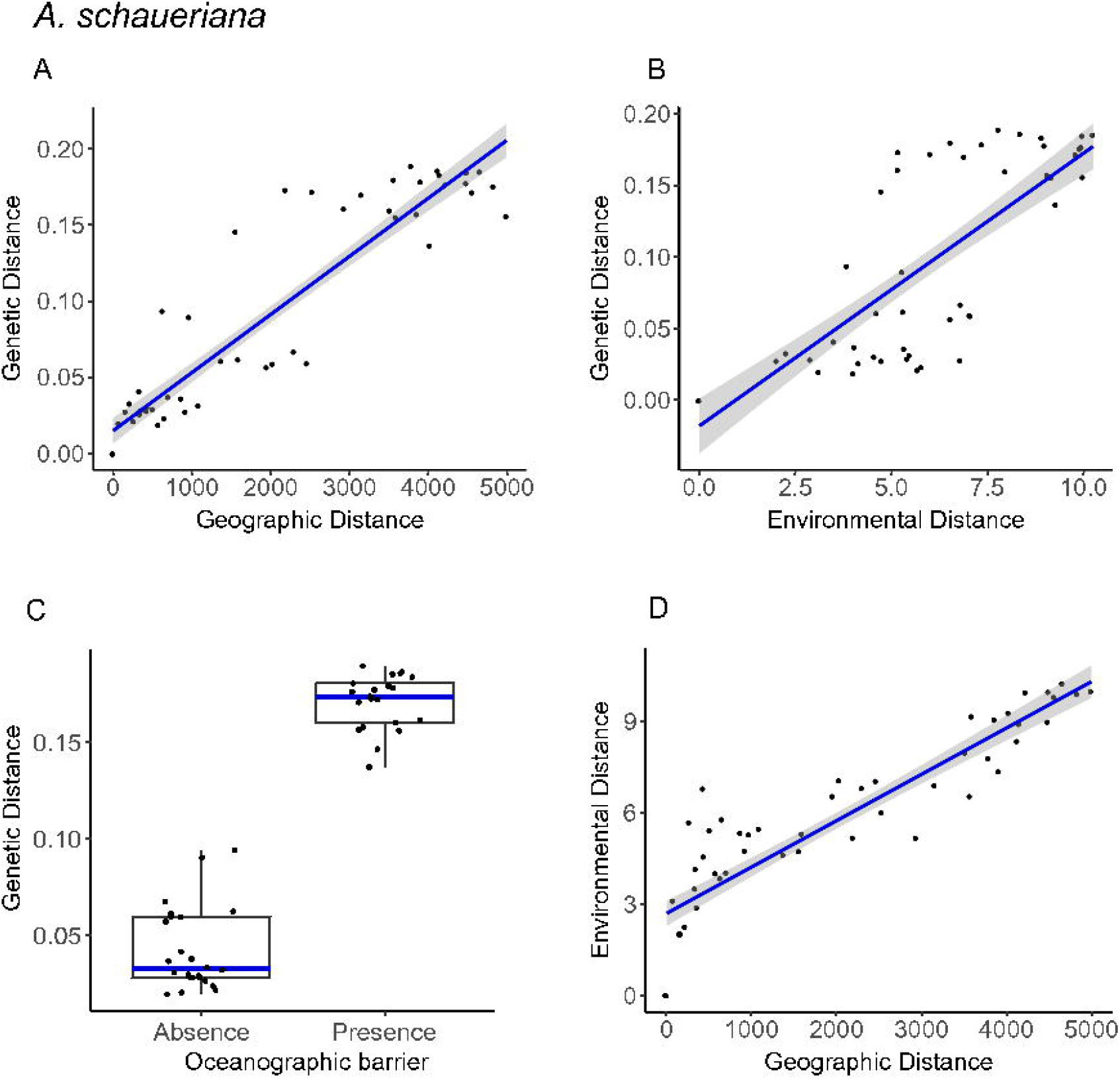
Graphical representations showing the correlations among genetic distance, geographical distance, environmental distance and the oceanographic barrier for *Avicennia schaueriana*. A, geographical distance vs. genetic distance. B, environmental distance vs. genetic distance. C, absence or presence of the oceanographic barrier between population pairs vs. genetic distance (boxplot). D, geographical distance vs. environmental distance. The relationships between genetic and geographical distances and between genetic distances and the oceanographic barrier were significant according to the partial Mantel tests (Table 2).

**Table 3.** Regression coefficient (β), coefficient of determination (*R²*) and significance (*P*) of the multiple matrix regression with randomization analysis in the association between genetic distance based on the total set of single nucleotide polymorphism molecular markers and combinations of geographical distance, environmental distance and the oceanographic barrier matrix of *Avicennia schaueriana* populations.

| Combination of landscape features | $\beta_{\text{geographical}}$ | $P$ | $\beta_{\text{environment}}$ | $P$ | $\beta_{\text{ocean barrier}}$ | $P$ | $R^2$ | $P$ |
| --- | --- | --- | --- | --- | --- | --- | --- | --- |
| D<sub>geo</sub> + D<sub>env</sub> | 0.13 | 0.001 | -0.48 | 0.01 | - | - | 0.86 | < 0.05 |
| D<sub>geo</sub> + M<sub>barrier</sub> | 0.25 | < 0.01 | - | - | 0.73 | < 0.01 | 0.94 | < 0.01 |
| D<sub>env</sub> + M<sub>barrier</sub> | - | - | 0.09 | 0.1 | 0.89 | < 0.01 | 0.93 | < 0.01 |
| D<sub>geo</sub> + D<sub>env</sub> + M<sub>barrier</sub> | 0.48 | < 0.01 | -0.18 | 0.14 | 0.67 | < 0.01 | 0.94 | 0.001 |

Abbreviations: D<sub>geo</sub>: Geographical distance; D<sub>env</sub>: Environmental distance; M<sub>barrier</sub>: Oceanographic barrier matrix.

For *A. germinans*, the simple Mantel test showed significant results only between the genetic distance and geographical distance (*r* = 0.78, *P* < 0.05) and between the genetic distance and environmental distance (*r* = 0.81, *P* = 0.01) based on the entire SNP dataset (Table 4 and Fig. 5). As observed for *A. schaueriana*, the geographical distance was also correlated with the environmental distance for this species (geographical vs. environmental distance: *r* = 0.86, *P* = 0.001; Fig. 5), implying that greater geographical distances correspond to greater environmental differences. However, the correlations between the oceanographic barrier matrix and the other two predictor variables were not significant (geographical distance vs. oceanographic barrier: *r* = 0.79, *P* = 0.16; environmental distance vs. oceanographic barrier: *r* = 0.78, *P* = 0.17). The partial Mantel tests showed significant values only for the environment, when conditioned on the geographical distance (*r* = 0.45, *P* = 0.01) (Table 4). In addition, the multivariate regression analysis of the combination of geographical and environmental distance showed significant values only for the environmental distance, which exerted an almost twofold greater effect than the geographical distance (β_environment_ = 0.55, *P* < 0.05, β_geographical_ = 0.28, *P* = 0.34) (Table 5). However, when we included the oceanographic barrier variable in the model, the regression coefficients for the three predictor variables were not significant (β_geographical_ = −0.07, *P* = 0.5; β_environment_ = 0.22, *P* = 0.2; β_ocean barrier_ = 0.84, *P* = 0.1) (Table 5). When we removed the samples from GT, which was the only sampling site south of the SEC, the genetic divergence was not correlated with either the geographical distance (partial Mantel: *r* = − 0.08, *P* = 0.61) or the environmental distance (partial Mantel: *r* = 0.17, *P* = 0.2; MMRR: β_geographical_ = −0.12, *P* = 0, 8; β_environment_ = 0.26, *P* = 0.6).

**Table 4.** Results of the simple and partial Mantel tests between genetic distance based on the total set of single nucleotide polymorphism molecular markers and the geographical distance, environmental distance and oceanographic barrier matrix for *Avicennia germinans* populations.

| Landscape feature | Controlled | $r$ | $P$ |
| --- | --- | --- | --- |
| Geographical distance |  | 0.78 | **0.04** |
| Environment distance |  | 0.81 | **0.01** |
| Oceanographic barrier |  | 0.95 | 0.16 |
| Geographical distance | Environment distance | 0.26 | 0.14 |
| Geographical distance | Oceanographic barrier | 0.14 | 0.25 |
| Environment distance | Geographical distance | 0.45 | **0.01** |
| Environment distance | Oceanographic barrier | 0.39 | 0.17 |
| Oceanographic barrier | Geographical distance | 0.88 | 0.12 |
| Oceanographic barrier | Environment distance | 0.88 | 0.15 |

Significant values are presented in bold.

**Figure 5.**
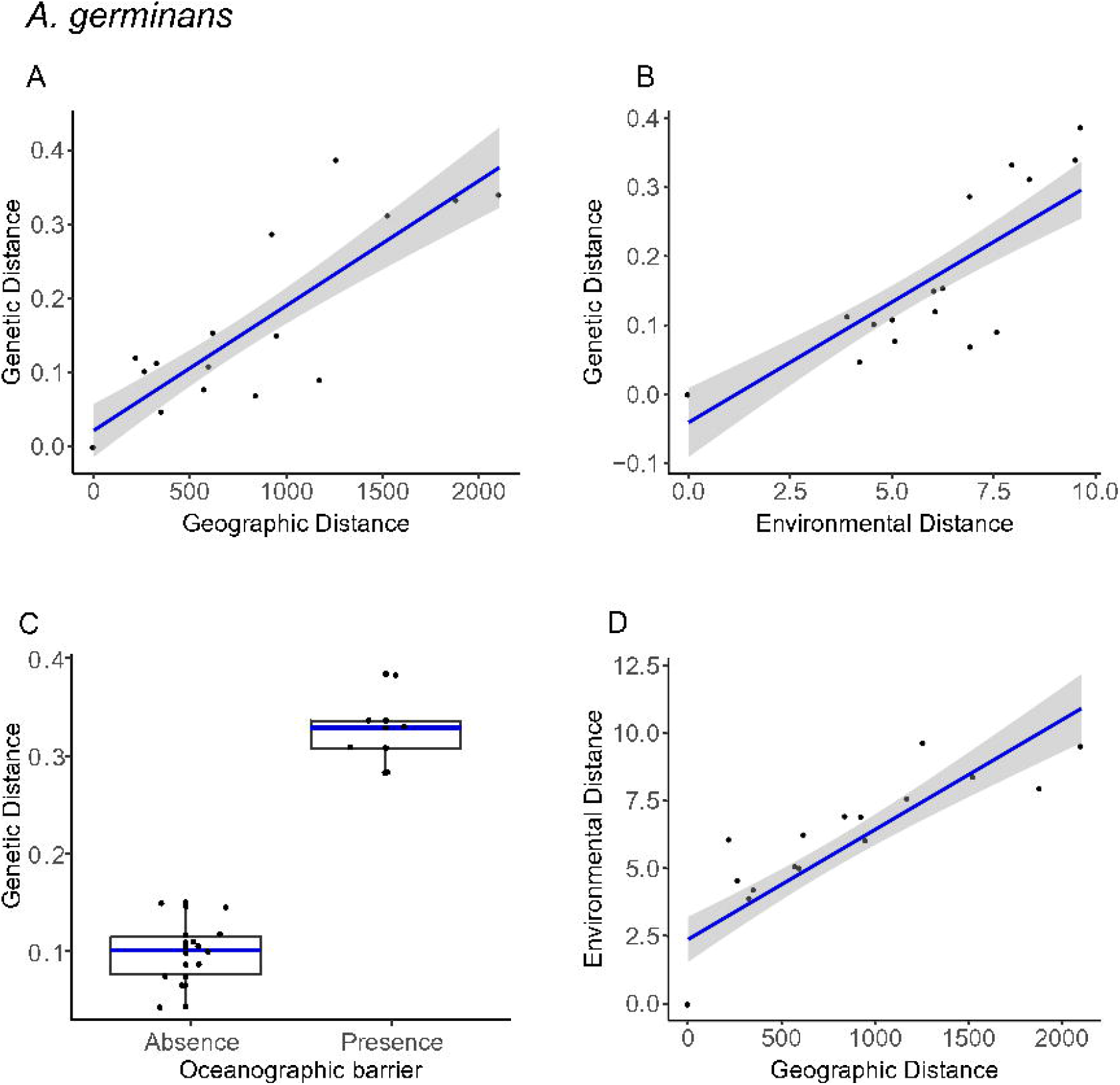
Graphical representations showing the correlations among genetic distance, geographical distance, environmental distance and the oceanographic barrier for *Avicennia germinans*. A, geographical distance vs. genetic distance. B, environmental distance vs. genetic distance. C, absence or presence of the oceanographic barrier between population pairs vs. genetic distance (boxplot). D, geographical distance vs. environmental distance. Among the relationships presented, only the environmental distance was significantly correlated with the genetic distance as indicated by the partial Mantel tests (Table 4).

**Table 5.** Regression coefficient (β), coefficient of determination (*R²*) and significance (*P*) of the multiple matrix regression with randomization analysis in the association between genetic distance based on the total set of single nucleotide polymorphism molecular markers and combinations of geographical distance, environmental distance and the oceanographic barrier matrix of *Avicennia germinans* populations.

| Combination of landscape feature | $\beta_{\text{geographical}}$ | $P$ | $\beta_{\text{environment}}$ | $P$ | $\beta_{\text{ocean barrier}}$ | $P$ | $R^2$ | $P$ |
| --- | --- | --- | --- | --- | --- | --- | --- | --- |
| D<sub>geo</sub> + D<sub>env</sub> | 0.28 | 0.34 | 0.55 | < 0.05 | - | - | 0.69 | < 0.05 |
| D<sub>geo</sub> + M<sub>barrier</sub> | 0.06 | 0.6 | - | - | 0.90 | 0.1 | 0.94 | 0.1 |
| D<sub>env</sub> + M<sub>barrier</sub> | - | - | 0.17 | 0.3 | 0.81 | 0.16 | 0.93 | < 0.01 |
| D<sub>geo</sub> + D<sub>env</sub> + M<sub>barrier</sub> | -0.07 | 0.5 | 0.22 | 0.2 | 0.84 | 0.1 | 0.94 | 0.06 |

Abbreviations: D<sub>geo</sub>: Geographical distance; D<sub>env</sub>: Environmental distance; M<sub>barrier</sub>: Oceanographic barrier matrix.

The results for genetic differentiation based on putative non-neutral SNPs showed the same patterns as those found for the total set of molecular markers for the two species. For *A. schaueriana*, both the geographical distance and the oceanographic barrier variable presented significant values based on this dataset (Supporting Information, Tables S11 and S12). For *A. germinans*, the genetic distance with evidence of natural selection was significantly correlated only with the environment conditioned on geography in partial Mantel tests (Supporting Information, Tables S13). Moreover, the combination of the environment and geography was the only significant model among the MMRR analyses (Supporting Information, Tables S14). Given the significant results for the environment for *A. germinans*, we analysed the correlation between each environmental variable and the genetic distance with evidence of natural selection. Significant correlations were observed for variables that vary along the latitudinal gradient of the species, such as the maximum solar radiation (partial Mantel: *r* = 0.95, *P* = 0.01; MMRR: β = 0.98, *P* = 0.01), precipitation in the warmest quarter (partial Mantel: *r* = 0.87, *P* = 0.05; MMRR: β = 0.95, *P* = 0.05), maximum temperature of the warmest month (partial Mantel: *r* = 0.83, *P* = 0.03; MMRR: β = 0.88, *P* < 0.05) and variations in the annual mean temperature (partial Mantel: *r* = 0.82, *P* = 0.01; MMRR: β = 0.88, *P* < 0.05) (Table 6, Supporting Information, Tables S15).

**Table 6.** Summary results of the partial Mantel test and multiple matrix regression with randomization (MMRR) analysis for on the associations between genetic distances based on the set of non-neutral molecular markers and the most important environmental variables for *Avicennia germinans*.

| Environmental variables | partial Mantel test |  | MMRR |  |
| --- | --- | --- | --- | --- |
| | $r$ | $P$-value | $\beta$ | $P$-value |
| Maximum solar radiation | 0.95 | 0.01 | 0.98 | 0.01 |
| Precipitation in the warmest quarter | 0.87 | 0.05 | 0.95 | 0.05 |
| Maximum temperature of the warmest month | 0.83 | < 0.05 | 0.88 | < 0.05 |
| Annual mean temperature | 0.82 | 0.01 | 0.88 | < 0.05 |

Partial Mantel tests were conditioned on geographical distance.

## DISCUSSION

The analysis of the geographical and environmental factors shaping neutral and adaptive genetic variation in heterogeneous environments is one of the main approaches used to understand the dynamics and evolutionary potential of natural populations (Schoville *et al.*, 2012; Li *et al.*, 2017). In the present study, we analysed the relative contributions of environmental and geographical distances and the presence of an oceanographic barrier to the genetic differentiation of populations of two dominant mangrove species in the Neotropics. We identified the relative importance of the main environmental variables that generate adaptation for one of these species, providing relevant information for decision-makers who will plan future efforts targeting conservation and the recovery of coastal vegetation in the face of increasing challenges resulting from anthropogenic, environmental and climate changes in this century.

Pairwise *F*_ST_ results revealed a variable degree of genetic divergence in both species, indicating the existence of substantially structured populations, particularly when samples to the north and south of the SEC are considered. These results corroborate the patterns of genetic structure reported in previous studies conducted with neutral molecular markers (Takayama *et al.*, 2008; Pil *et al.*, 2011; Mori *et al.*, 2015; Francisco *et al.*, 2018) and indicate that regardless of the characteristics of these markers with high (microsatellite; Vieira *et al.*, 2016) or low (SNPs; Morin *et al.*, 2004) mutation rates, the evolutionary processes that led to this divergence must be intense or ancient.

The geographical distances between sampling sites contributed significantly to the genetic divergence of *A. schaueriana*, suggesting that spatial distance plays a fundamental role in the genetic divergence of populations of this species. This model appears to be common in studies of plants in general (Sexton *et al.*, 2014; Segarra-Moragues *et al.*, 2016; Cruz-Nicolás *et al.*, 2019) and mangroves in particular (Cerón-Souza *et al.*, 2010; Sandoval-Castro *et al.*, 2014; Kennedy *et al.*, 2016; Binks *et al.*, 2019; Ochoa-Zavala *et al.*, 2019). Although evidence of water dispersion over long distances exists for mangrove species (Nettel & Dodd, 2007; Takayama *et al.*, 2013; Mori *et al.*, 2015; Van der Stocken *et al.*, 2019b), our results indicate that the large geographical extent and major oceanic currents of the Brazilian coast physically limit the dispersal of *Avicennia* species.

For *A. germinans* populations, when the geographical distance was controlled by another covariate (Table 4) it did not show significant correlations for the total set of samples or for the five sampling locations north of the SEC, which were distributed in a fairly geographically continuous habitat. We hypothesized that the genetic differentiation ranging from 0.047 to 0.387 observed among sampling locations might result from an IBB effect caused by the SEC acting as a barrier to the dispersal of propagules. However, we did not find a significant correlation between genetic differentiation and the presence of the SEC. Although we did not find significant results, the explained variation and the correlation coefficient for the IBB model were higher than that of the other two models for this species (Tables 4 and 5). In addition, we also observed a slight separation in the point clouds in the scatter plots of both species between the genetic and geographical distances and between the genetic and environmental distances (Figs 4, 5), which seem to reflect the influence of the SEC bifurcation on genetic differentiation among populations. The non-significant values probably reflected insufficient sampling south of the SEC, where only a single location, GT, was used. In this context, we suggest that future efforts should address the limitations of our study to generate more conclusive IBB results for *A. germinans*.

In contrast, we obtained statistical evidence for the action of the SEC as a barrier to gene flow in *A. schaueriana*. Our results suggest that IBB is one of the main models for genetic differentiation among populations of this species. This model has also been shown in populations of *A. germinans* and *R. mangle* in Central America, whose patterns of genetic diversity were consistent with the patterns of ocean circulation in the east tropical Pacific (Cerón-Souza *et al.*, 2015). Our results corroborate the findings reported by Mori *et al.* (2015), who suggested that the neutral genetic divergence observed for *A. schaueriana* might have been shaped by marine currents. Based on our results, the bifurcated flow of marine currents along the Atlantic coast of South America might play a key role as a driver of the genetic differentiation observed in other species of mangrove or species associated with this ecosystem, such as *R. mangle* (Pil *et al.*, 2011; Francisco *et al.*, 2018) and *H. pernambucensis* (Takayama *et al.*, 2008). Our findings showed statistically that the SEC is an important driver of the genetic structure of mangrove species; however, coastal and ocean currents vary temporally in strength and directionality (Van der Stocken *et al.*, 2019a). For example, the SEC splits into the BC and the NBC, which have different speeds and directions (Fig. 1; Lumpkin & Johnson, 2013). The NBC is faster than the BC, favouring the spread of propagules from south to north, as observed in previous studies (Mori *et al.*, 2015; Francisco *et al.*, 2018). Additionally, the direction of flow of coastal currents might influence the direction of gene flow among populations. Therefore, future investigations of the dynamics of these currents, not as a static barrier but including different levels of resistance to gene flow, might provide more realistic insights into their effects on the distribution of the genetic variation of species dispersed by sea currents.

We also identified an IBE pattern in the structure of the genetic diversity of *A. germinans*. For this species, this model presented significant values in relationship to neutral processes (Tables 4 and 5; Supporting Information, Tables S13 and S14), suggesting an important role of environmental heterogeneity in the reproduction and survival of migrant individuals. Many species showed this same pattern with the IBE model (Mitchell-Olds *et al.*, 2007; Byars *et al.*, 2009; Barker *et al.*, 2011; Vernesi *et al.*, 2012; Shafer & Wolf, 2013; Dennenmoser *et al.*, 2014; Sexton *et al.*, 2014; Manthey & Moyle, 2015; Rodríguez-Zárate *et al.*, 2018; Jiang *et al.*, 2019), indicating that environmental heterogeneity might be the main factor underlying the geographical distance (Shafer & Wolf, 2013; Sexton *et al.*, 2014; Beheregaray *et al.*, 2015).

We also identified the key environmental factors underlying the organization of the genetic diversity of *A. germinans*. The genetic variation with evidence of selection was explained mainly by atmospheric temperature patterns, precipitation regimens and solar radiation (Table 6; Supporting Information, Table S15). These results corroborate the findings reported by Cruz *et al.* (2020), who performed a functional characterization of putative loci under selection and suggested that differential precipitation regimens play a fundamental role in genetic divergence between populations of this species. Additionally, our findings support the results reported by Bajay *et al.* (2018), who found different profiles of gene expression among populations of *R. mangle* located at contrasting latitudes of the Brazilian coast, with differentially expressed genes putatively involved in the responses to variations in temperature, solar radiation and precipitation. The results obtained for *A. germinans* corroborate the data from other studies reporting that precipitation and temperature variables are limiting factors regulating the distribution of coastal species (McKee *et al.*, 2012; Soares *et al.*, 2012; Osland *et al.*, 2016; Duke *et al.*, 2017; Cavanaugh *et al.*, 2018; Ximenes *et al.*, 2018). Although we acknowledge the need for complementary studies that avoid molecular spandrels (Barrett & Hoekstra, 2011), our findings, together with the results published by Cruz *et al.* (2020), provide new evidence for the role of local adaptation in the distribution of the genetic diversity of *A. germinans*.

Our results are particularly relevant in light of the climate changes that have been occurring in the last few decades. With them, further investigations on the responses of *A. germinans* to future changes in predicted increases in the average annual temperature and rainfall regimens (IPCC, 2014) can be directed. These results also have implications for conservation management and planning decisions (Friess *et al.*, 2019). With a better understanding of the role of environmental variables in modelling the genetic variation of *A. germinans* on the South American coast, our results contribute to the definition of evolutionarily significant units for this species, thus maintaining its evolutionary potential (Fraser & Bernatchez, 2001). In addition, based on the results showing the remarkable pattern of genetic divergence of *A. schaueriana*, which is attributable, in part, to the action of the current of the marine surface studied here, we suggest the consideration of genetic banks north and south from the SEC as conservation measures for the maintenance of biodiversity.

## CONCLUSIONS

Our results provide new evidence that the combined actions of the geographical distance, ocean currents and environmental gradient contribute to the evolution of spatial genetic divergence within two Neotropical *Avicennia* species growing on the Atlantic coast of South America. Our findings reveal that geographical distance and ocean currents can influence the pattern of gene flow in *A. schaueriana* populations, with a greater predictive ability found for IBB. We found an IBE pattern in *A. germinans* and evidence of adaptations to climate variables that have changed rapidly in recent decades. In this context, our results provide a basis for understanding the role of geographical and environmental factors in shaping genetic variation in *Avicennia* species of the South American coast. These results contribute to our understanding of the evolution of genetic diversity within this genus and provide information that might be relevant to other coastal organisms. Additionally, our findings will guide further research on the responses of these species to climate change and contribute to effective conservation plans for mangroves and other species with similar mechanisms of dispersion.

## Supporting information

Eletronic suplementary material Tables

## ACKNOWLEDGEMENTS

We thank Alessandro Alves Pereira and Prianda Rios Laborda, in addition to three anonymous reviewers, for critically reading the manuscript. We are also truly grateful for the financial support from the Coordination for the Improvement of Higher Education Personnel (CAPES Computational Biology Program), the São Paulo Research Foundation (FAPESP) and the Brazilian National Council for Scientific and Technological Development (CNPq). M.F.S. received fellowships from FAPESP (grant numbers 2018/18431-1 and 2020/00203-2) and the CAPES Computational Biology Program (grant number 88881.185134/2018-01). M.V.C. received a fellowship from FAPESP (grant number 2013/26793-7). J.D.V. received a fellowship from CNPq (grant number 153973/2018-8). G.M.M. received fellowships from FAPESP (grant numbers 2013/08086-1 and 2014/22821-9) and a research award from CNPq (grant number 448286/2014-9). A.P.S. received fellowships from the CAPES Computational Biology Program (grant number 88882.160095/2013-01) and a research fellowship from CNPq (grant number 312777/2018-3). The authors have no competing interests to declare.

## SUPPORTING INFORMATION

Additional Supporting Information may be found in the online version of this article at the publisher’s web-site:

**Figure S1.** Groups (red boxes) of correlated environmental variables retrieved from the public data platforms WorldClim (Fick & Hijmans, 2017), Marspec (Sbrocco & Barber, 2013), and ECDS (Klein *et al.*, 2013). The cutoff value for Pearson’s correlation coefficient was set to 0.8. *Environmental variables retained for subsequent analysis.

**Table S1.** Environmental data matrix for the ten sampling points of *Avicennia schaueriana*. Sample codes are denoted as in Table 1.

**Table S2.** Environmental data matrix for the six sampling points of *Avicennia germinans*. Sample codes are denoted as in Table 1.

**Table S3.** Interpopulation genetic differentiation based on the set of markers with evidence of natural selection estimated by *F*_ST_ (Wright) pairs for the *Avicennia schaueriana* collection. Sample codes are denoted as in Table 1.

**Table S4.** Interpopulation genetic differentiation based on the set of markers with evidence of natural selection estimated by *F*_ST_ (Wright) pairs for the *Avicennia germinans* collection. Sample codes are denoted as in Table 1.

**Table S5**. Pairwise geographical distances for *Avicennia schaueriana*. Sample codes are denoted as in Table 1.

**Table S6.** Pairwise geographical distances for *Avicennia germinans*. Sample codes are denoted as in Table 1.

**Table S7.** Pairwise environmental distances based on each environmental variable for *Avicennia schaueriana*. Sample codes are denoted as in Table 1.

**Table S8.** Pairwise environmental distances based on each environmental variable for *Avicennia germinans.* Sample codes are denoted as in Table 1.

**Table S9.** Pairwise environmental distances for *Avicennia schaueriana*. Sample codes are denoted as in Table 1.

**Table S10.** Pairwise environmental distances for *Avicennia germinans*. Sample codes are denoted as in Table 1.

**Table S11.** Results of the simple and partial Mantel tests between genetic distance based on the set of non-neutral molecular markers and the geographical distance, environmental distance and oceanographic barrier matrix for *Avicennia schaueriana* populations.

**Table S12.** Regression coefficient (β), coefficient of determination (*R²*) and significance (*P*) of the multiple matrix regression with randomization (MMRR) analysis in the association between genetic distance based on the set of non-neutral molecular markers and combinations between geographical distance, environmental distance and the oceanographic barrier matrix of *Avicennia schaueriana* populations.

**Table S13.** Results of the simple and partial Mantel tests between genetic distance based on the set of non-neutral molecular markers and the geographical distance, environmental distance and oceanographic barrier matrix for *Avicennia germinans* populations.

**Table S14.** Regression coefficient (β), coefficient of determination (*R²*) and significance (*P*) of the multiple matrix regression with randomization (MMRR) analysis in the association between genetic distance based on the set of non-neutral molecular markers and combinations of geographical distance, environmental distance and the oceanographic barrier matrix of *Avicennia germinans* populations.

**Table S15.** Results of the partial Mantel test and multiple matrix regression with randomization (MMRR) analysis for the associations between genetic distance based on the set of non-neutral molecular markers and the environmental variables for *Avicennia germinans*.

## SHARED DATA

*Avicennia germinans* SNP genotype data are available from the Dryad Digital Repository (Cruz *et al*., 2019). *A. schaueriana* SNP genotype data are available from the article: doi:10.1038/s41598-019-56469-w (supplementary datasets 1 and 2).

## REFERENCES

Antao T, Lopes A, Lopes RJ, Beja-Pereira A, Luikart G. 2008. LOSITAN: a workbench to detect molecular adaptation based on a F st -outlier method. BMC Bioinformatics 9: 323.

Bajay SK, Cruz MV, da Silva CC, Murad NF, Brandão MM, de Souza AP. 2018. Extremophiles as a model of a natural ecosystem: transcriptional coordination of genes reveals distinct selective responses of plants under climate change scenarios. Frontiers in Plant Science 9: 1376.

Barker JSF, Frydenberg J, Sarup P, Loeschcke V. 2011. Altitudinal and seasonal variation in microsatellite allele frequencies of *Drosophila buzzatii*. Journal of Evolutionary Biology 24: 430–439.

Barrett RD, Hoekstra HE. 2011. Molecular spandrels: tests of adaptation at the genetic level. Nature Reviews Genetics 12: 767–780.

Barton NH. 1979. Gene flow past a cline. Heredity 43: 333–339.

Beheregaray LB, Cooke GM, Chao NL, Landguth EL. 2015. Ecological speciation in the tropics: insights from comparative genetic studies in Amazonia. Frontiers in Genetics 5: 477.

Binks RM, Byrne M, McMahon K, Pitt G, Murray K, Evans RD. 2019. Habitat discontinuities form strong barriers to gene flow among mangrove populations, despite the capacity for long-distance dispersal. Diversity and Distributions 25: 298–309.

Bradburd GS, Ralph PL, Coop GM. 2013. Disentangling the effects of geographic and ecological isolation on genetic differentiation. Evolution 67: 3258–3273.

Byars SG, Parsons Y, Hoffmann AA. 2009. Effect of altitude on the genetic structure of an Alpine grass, Poa hiemata. Annals of Botany 103: 885–899.

Cavanaugh KC, Osland MJ, Bardou R, Hinojosa-Arango G, López-Vivas JM, Parker JD, Rovai AS. 2018. Sensitivity of mangrove range limits to climate variability. Global Ecology and Biogeography 27: 925–935.

Cerón-Souza I, Rivera-Ocasio E, Medina E, Jiménez JA, McMillan WO, Bermingham E. 2010. Hybridization and introgression in New World red mangroves, *Rhizophora* (Rhizophoraceae). American Journal of Botany 97: 945–957.

Cerón-Souza I, Gonzalez EG, Schwarzbach AE, Salas-Leiva DE, Rivera-Ocasio E, Toro-Perea N, Bermingham E, McMillan WO. 2015. Contrasting demographic history and gene flow patterns of two mangrove species on either side of the Central American Isthmus. Ecology and Evolution 5: 3486–3499

Chapman AD. 2005. Principles and methods of data cleaning – primary species and species occurrence data, version 1. 0. Copenhagen: Report for the Global Biodiversity Information Facility. Available online at http://www.gbif.org/document/80528.

Cruz MV, Mori GM, Oh DH, Dassanayake M, Zucchi MI, Oliveira RS, Souza AP. 2020. Molecular responses to freshwater limitation in the mangrove tree *Avicennia germinans* (Acanthaceae). Molecular Ecology 29: 344–362.

Cruz MV, Mori GM, Signori-Müller C, da Silva CC, Oh D-H, Dassanayake M, Zucchi MI, Oliveira RS, de Souza AP. 2019. Local adaptation of a dominant coastal tree to freshwater availability and solar radiation suggested by genomic and ecophysiological approaches. Scientific Reports 9: 27.

Cruz-Nicolás J, Giles-Pérez G, González-Linares E, Múgica-Gallart J, Lira-Noriega A, Gernandt DS, Eguiarte LE, Jaramillo-Correa JP. 2019. Contrasting evolutionary processes drive morphological and genetic differentiation in a subtropical fir (*Abies*, Pinaceae) species complex. Botanical Journal of the Linnean Society 192: 401–420.

Cushman SA, McKelvey KS, Hayden J & Schwartz MK. 2006. Gene flow in complex landscapes: Testing multiple hypotheses with causal modeling. American Naturalist 168: 486–499.

Danecek P, Auton A, Abecasis G, Albers CA, Banks E, DePristo MA, Handsaker RE, Lunter G, Marth GT, Sherry ST, McVean G, Durbin R. 2011. The variant call format and VCF tools. Bioinformatics 27: 2156–2158.

Dennenmoser S, Rogers SM, Vamosi SM. 2014. Genetic population structure in prickly sculpin (*Cottus asper*) reflects isolation-by-environment between two life-history ecotypes. Biological Journal of the Linnean Society 113: 943–957.

Duke NC. 1991. A systematic revision of the mangrove genus Avicennia (Avicenniaceae) in Australasia. Australian Systematic Botany 4: 299.

Duke NC, Kovacs JM, Griffiths AD, Preece L, Hill DJE, van Oosterzee P, Mackenzie J, Morning HS, Burrows D. 2017. Large-scale dieback of mangroves in Australia. Marine and Freshwater Research 68: 1816.

Fick SE, Hijmans RJ. 2017. WorldClim 2: new 1-km spatial resolution climate surfaces for global land areas. International Journal of Climatology 37: 4302–4315.

Francisco PM, Mori GM, Alves FM, Tambarussi EV, de Souza AP. 2018. Population genetic structure, introgression, and hybridization in the genus Rhizophora along the Brazilian coast. Ecology and Evolution 8: 3491–3504.

Fraser DJ, Bernatchez L. 2001. Adaptive evolutionary conservation: towards a unified concept for defining conservation units. Molecular Ecology 10: 2741–2752.

Frichot E, François O. 2015. LEA: an R package for landscape and ecological association studies. Methods in Ecology and Evolution 6: 925–929.

Friess D, Rogers K, Lovelock C, Krauss K, Hamilton S, Lee S, Lucas R, Primavera J, Rajkaran A, Shi S. 2019. The state of the world’s mangrove forests: past, present, and future. Annual Review of Environment and Resources 44: 1–27.

Giri C, Ochieng E, Tieszen LL, Zhu Z, Singh A, Loveland T, Masek J, Duke N. 2011. Status and distribution of mangrove forests of the world using earth observation satellite data. Global Ecology and Biogeography 20: 154–159.

Goslee SC & Urban DL. 2007. The ecodist package for dissimilarity-based analysis of ecological data. Journal of Statistical Software 22: 1–19.

Goudet J. 2005. hierfstat, a package for r to compute and test hierarchical F-statistics. Molecular Ecology Notes 5: 184–186.

Hamilton S.E. 2020. Botany of Mangroves. In: Mangroves and Aquaculture. Coastal Research Library, vol 33. Springer, Cham. https://doi.org/10.1007/978-3-030-22240-6_1.

Hijmans RJ. 2017. raster: geographic data analysis and modelling. R package version 2.6–7. Available at: https://cran.r-project.org/package=raster.

Holderegger R, Buehler D, Gugerli F, Manel S. 2010. Landscape genetics of plants. Trends in Plant Science 15: 675–683.

IPCC, 2014: Climate Change 2014: Synthesis Report. Contribution of Working Groups I, II and III to the Fifth Assessment Report of the Intergovernmental Panel on Climate Change [Core Writing Team, R.K. Pachauri and L.A. Meyer (eds.)]. IPCC, Geneva, Switzerland, 151 pp.

Jasechko S, Sharp ZD, Gibson JJ, Birks SJ, Yi Y, Fawcett PJ. 2013. Terrestrial water fluxes dominated by transpiration. Nature 496: 347–350.

Jiang S, Luo MX, Gao RH, Zhang W, Yang YZ, Li YJ, Liao PC. 2019. Isolation-by-environment as a driver of genetic differentiation among populations of the only broad-leaved evergreen shrub *Ammopiptanthus mongolicus* in Asian temperate deserts. Scientific Reports 9: 12008.

Joost S, Bonin A, Bruford MW, Després L, Conord C, Erhardt G, Taberlet P. 2007. A Spatial Analysis Method (SAM) to detect candidate loci for selection: towards a landscape genomics approach to adaptation. Molecular Ecology 16: 3955–3969.

Kennedy JP, Pil MW, Proffitt CE, Boeger WA, Stanford AM, Devlin DJ. 2016. Postglacial expansion pathways of red mangrove, Rhizophora mangle, in the Caribbean Basin and Florida. American Journal of Botany 103: 260–276.

Klein T, Langner J, Frankenberg B, Svensson J, Broman B, Bennet C, Langborg T. 2013. ECDS - a Swedish research infrastructure for the open sharing of environment and climate data. Data Science Journal 12: 1–9.

Kovach RP, Gharrett AJ, Tallmon DA. 2012. Genetic change for earlier migration timing in a pink salmon population. Proceedings of the Royal Society B: Biological Sciences 279: 3870–3878.

Lee CR, Mitchell-Olds T. 2011. Quantifying effects of environmental and geographical factors on patterns of genetic differentiation. Molecular Ecology 20: 4631–4642.

Legendre P. 1993. Spatial Autocorrelation: Trouble or New Paradigm? Ecology 74:1659–1673.

Legendre, P., and L. Legendre. 2012. Numerical ecology, 3rd edn. Elsevier, Amsterdam.

Leroy B, Meynard CN, Bellard C, Courchamp F. 2016. virtualspecies, an R package to generate virtual species distributions. Ecography 39: 599–607.

Li X, Duke NC, Yang Y, Huang L, Zhu Y, Zhang Z, Zhou R, Zhong C, Huang Y, Shi S. 2016. Re-evaluation of phylogenetic relationships among species of the Mangrove Genus Avicennia from Indo-West Pacific based on multilocus analyses. PLoS One 11: e0164453.

Li Y, Zhang XX, Mao RL, Yang J, Miao CY, Li Z, Qiu YX. 2017. Ten years of landscape genomics: challenges and opportunities. Frontiers in Plant Science 8: 2136.

Lowry DB. 2010. Landscape evolutionary genomics. Biology Letters 6: 502–504.

Lumpkin R, Johnson GC. 2013. Global ocean surface velocities from drifters: mean, variance, El Niño-Southern Oscillation response, and seasonal cycle. Journal of Geophysical Research: Oceans 118: 2992–3006.

Luu K, Bazin E, Blum MGB. 2017. pcadapt: anRpackage to perform genome scans for selection based on principal component analysis. Molecular Ecology Resources 17: 67–77.

Mannion PD, Upchurch P, Benson RBJ, Goswami A. 2014. The latitudinal biodiversity gradient through deep time. Trends in Ecology & Evolution 29: 42–50.

Manthey JD, Moyle RG. 2015. Isolation by environment in White-breasted Nuthatches (*Sitta carolinensis*) of the Madrean Archipelago sky islands: a landscape genomics approach. Molecular Ecology 24: 3628–3638.

McKee K, Rogers K, Saintilan N. 2012. Response of salt marsh and Mangrove wetlands to changes in atmospheric CO2, climate, and sea level. In: Middleton BA, ed. Global change and the function and distribution of wetlands. Dordrecht, Netherlands: Springer, 63–96.

Middleton BA. 2012. Global change and the function and distribution of wetlands. Dordrecht, Netherlands: Springer.

Mitchell-Olds T, Willis JH, Goldstein DB. 2007. Which evolutionary processes influence natural genetic variation for phenotypic traits? Nature Reviews Genetics 8: 845–856.

Mori GM, Zucchi MI, Souza AP. 2015. Multiple-geographic-scale genetic structure of two Mangrove tree species: the roles of mating system, hybridization, limited dispersal and extrinsic factors. PLoS One 10: e0118710.

Morin PA, Luikart G, Wayne RK, The SNP Workshop Group. 2004. SNPs in ecology, evolution and conservation. Trends in Ecology & Evolution 19: 208–216.

Morrisey DJ, Swales A, Dittmann S, Morrison MA, Lovelock CE, Beard CM. 2010. The ecology and management of temperate mangroves. In: Gibson RN, Atkinson RJA, Gordon JDM, eds. Oceanography and marine biology: an annual review. Boca Raton, FL: Chapman and Hall/CRC, 43–160.

Muñoz NJ, Farrell AP, Heath JW, Neff BD. 2015. Adaptive potential of a Pacific salmon challenged by climate change. Nature Climate Change 5: 163–166.

Murray KD, Janes JK, Jones A, Bothwell HM, Andrew RL, Borevitz JO. 2019. Landscape drivers of genomic diversity and divergence in woodland Eucalyptus. Molecular Ecology 28: 5232–5247.

Nettel A, Dodd RS. 2007. Drifting propagules and receding swamps: genetic footprints of mangrove recolonization and dispersal along tropical coasts. Evolution 61: 958–971.

Ochoa-Zavala M, Jaramillo-Correa JP, Piñero D, Nettel-Hernanz A, Núñez-Farfán J. 2019. Contrasting colonization patterns of black mangrove (*Avicennia germinans* (L.) L.) gene pools along the Mexican coasts. Journal of Biogeography 46: 884–898.

Osland MJ, Enwright NM, Day RH, Gabler CA, Stagg CL, Grace JB. 2016. Beyond just sea-level rise: considering macroclimatic drivers within coastal wetland vulnerability assessments to climate change. Global Change Biology 22: 1–11.

Osland MJ, Feher LC, Griffith KT, Cavanaugh KC, Enwright NM, Day RH, Stagg CL, Krauss KW, Howard RJ, Grace JB, Rogers K. 2017. Climatic controls on the global distribution, abundance, and species richness of mangrove forests. Ecological Monographs 87: 341–359.

Pil MW, Boeger MRT, Muschner VC, Pie MR, Ostrensky A, Boeger WA. 2011. Postglacial north-south expansion of populations of *Rhizophora mangle* (Rhizophoraceae) along the Brazilian coast revealed by microsatellite analysis. American Journal of Botany 98: 1031–1039.

R Core Team (2019). R: A language and environment for statistical computing. R Foundation for Statistical Computing, Vienna, Austria. URL https://www.R-project.org/.

Robertson JM, Duryea MC & Zamudio KR. 2009. Discordant patterns of evolutionary differentiation in two Neotropical treefrogs. Molecular Ecology 18: 1375–1395.

Rodríguez-Zárate CJ, Sandoval-Castillo J, van Sebille E, Keane RG, Rocha-Olivares A, Urteaga J, Beheregaray LB. 2018. Isolation by environment in the highly mobile olive ridley turtle (*Lepidochelys olivacea*) in the eastern Pacific. Proceedings of the Royal Society B: Biological Sciences 285: 20180264.

Saintilan N, Khan NS, Ashe E, Kelleway JJ, Rogers K, Woodroffe CD, Horton BP. 2020. Thresholds of mangrove survival under rapid sea level rise. Science 368: 1118.

Sandoval-Castro E, Dodd RS, Riosmena-Rodríguez R, Enríquez-Paredes LM, Tovilla-Hernández C, López-Vivas JM, Aguilar-May B, Muñiz-Salazar R. 2014. Post-glacial expansion and population genetic divergence of Mangrove Species *Avicennia germinans* (L.) stearn and *Rhizophora mangle* L. along the Mexican Coast. PLoS One 9: e93358.

Savolainen O, Pyhäjärvi T, Knürr T. 2007. Gene flow and local adaptation in trees. Annual Review of Ecology, Evolution, and Systematics 38: 595–619.

Sbrocco EJ, Barber PH. 2013. MARSPEC: ocean climate layers for marine spatial ecology. Ecology 94: 979.

Schaeffer-Novelli Y, Cintrón-Molero G, Adaime RR, de Camargo TM, Cintron-Molero G, de Camargo TM. 1990. Variability of Mangrove ecosystems along the Brazilian Coast. Estuaries 13: 204–218.

Schoville SD, Bonin A, François O, Lobreaux S, Melodelima C, Manel S. 2012. Adaptive genetic variation on the landscape: methods and cases. Annual Review of Ecology, Evolution, and Systematics 43: 23–43.

Segarra-Moragues JG, Carrión Marco Y, Castellanos MC, Molina MJ, García-Fayos P. 2016. Ecological and historical determinants of population genetic structure and diversity in the Mediterranean shrub *Rosmarinus officinalis* (Lamiaceae). Botanical Journal of the Linnean Society 180: 50–63.

Sexton JP, Hangartner SB, Hoffmann AA. 2014. Genetic isolation by environment or distance: which pattern of gene flow is most common? Evolution 68: 1–15.

Shafer ABA, Wolf JBW. 2013. Widespread evidence for incipient ecological speciation: a meta-analysis of isolation-by-ecology. Ecology Letters 16: 940–950.

Smouse PE, Long JC & Sokal RR. 2012. Multiple regression and correlation extensions of the Mantel test of matrix correspondence. Systematic Zoology. 35: 627–632.

Soares MLG, Estrada GCD, Fernandez V, Tognella MMP. 2012. Southern limit of the Western South Atlantic Mangroves: assessment of the potential effects of global warming from a biogeographical perspective. Estuarine, Coastal and Shelf Science 101: 44–53.

Sork VL. 2016. Gene flow and natural selection shape spatial patterns of genes in tree populations: implications for evolutionary processes and applications. Evolutionary Applications 9: 291–310.

Spalding M, Blasco F, Field CD. 1997. World mangrove atlas. Japan, Okinawa: The International Society for Mangroves Ecosystems.

Storfer A, Murphy MA, Spear SF, Holderegger R, Waits LP. 2010. Landscape genetics: where are we now? Molecular Ecology 19: 3496–3514.

Storfer A, Patton A, Fraik AK. 2018. Navigating the interface between landscape genetics and landscape genomics. Frontiers in Genetics 9: 68.

Takayama K, Tamura M, Tateishi Y, Webb EL, Kajita T. 2013. Strong genetic structure over the American continents and transoceanic dispersal in the mangrove genus *Rhizophora* (Rhizophoraceae) revealed by broad-scale nuclear and chloroplast DNA analysis. American Journal of Botany 100: 1191–1201.

Takayama K, Tateishi Y, Murata JIN, Kajita T. 2008. Gene flow and population subdivision in a pantropical plant with sea-drifted seeds *Hibiscus tiliaceus* and its allied species: evidence from microsatellite analyses. Molecular Ecology 17: 2730–2742.

Tomlinson. 1986. The Botany of Mangroves. New York: Cambridge University Press.

Van der Stocken T, Carroll D, Menemenlis D, Simard M, Koedam N. 2019a. Global-scale dispersal and connectivity in mangroves. Proceedings of the National Academy of Sciences 116: 915–922.

Van der Stocken T, Wee AKS, De Ryck DJR, Vanschoenwinkel B, Friess DA, Dahdouh-Guebas F, Simard M, Koedam N, Webb EL. 2019b. A general framework for propagule dispersal in mangroves. Biological Reviews 94: 1547–1575.

Vernesi C, Rocchini D, Pecchioli E, Neteler M, Vendramin GG, Paffetti D. 2012. A landscape genetics approach reveals ecological-based differentiation in populations of holm oak (*Quercus ilex* L.) at the northern limit of its range. Biological Journal of the Linnean Society 107: 458–467.

Vieira MLC, Santini L, Diniz AL, Munhoz CF. 2016. Microsatellite markers: what they mean and why they are so useful. Genetics and Molecular Biology 39: 312–328.

Vincent B, Dionne M, Kent MP, Lien S, Bernatchez L. 2013. Landscape genomics in *Atlantic salmon* (Salmo Salar): searching for gene-environment interactions driving local adaptation. Evolution 67: 3469–3487.

Wang IJ. 2013. Examining the full effects of landscape heterogeneity on spatial genetic variation: a multiple matrix regression approach for quantifying geographic and ecological isolation. Evolution 67: 3403–3411.

Wang IJ, Bradburd GS. 2014. Isolation by environment. Molecular Ecology 23: 5649–5662.

Wang IJ, Glor RE, Losos JB. 2013. Quantifying the roles of ecology and geography in spatial genetic divergence. Ecology Letters 16: 175–182.

Wee AKS, Mori GM, Lira CF, Núñez-Farfán J, Takayama K, Faulks L, Shi S, Tsuda Y, Suyama Y, Yamamoto T, Iwasaki T, Nagano Y, Wang Z, Watanabe S, Kajita T. 2019. The integration and application of genomic information in Mangrove conservation. Conservation Biology 33: 206–209.

Wright S. 1943. Isolation by distance. Genetics 28: 114–138.

Wright S. 1949. The genetical structure of populations. Annals of Eugenics 15: 323–354.

Wu Z, Yu D, Li X & Xu X. 2016. Influence of geography and environment on patterns of genetic differentiation in a widespread submerged macrophyte, Eurasian watermilfoil (Myriophyllum spicatum L., Haloragaceae). Ecology and Evolution 6: 460–468.

Ximenes A, Ponsoni L, Lira C, Koedam N, Dahdouh-Guebas F. 2018. Does sea surface temperature contribute to determining range limits and expansion of Mangroves in Eastern South America (Brazil)? Remote Sensing 10: 1787.

