## Supplementary information Figure S1 for "Geographical and environmental contributions to genomic divergence in mangrove forests"

**Electronic Supplementary Material**

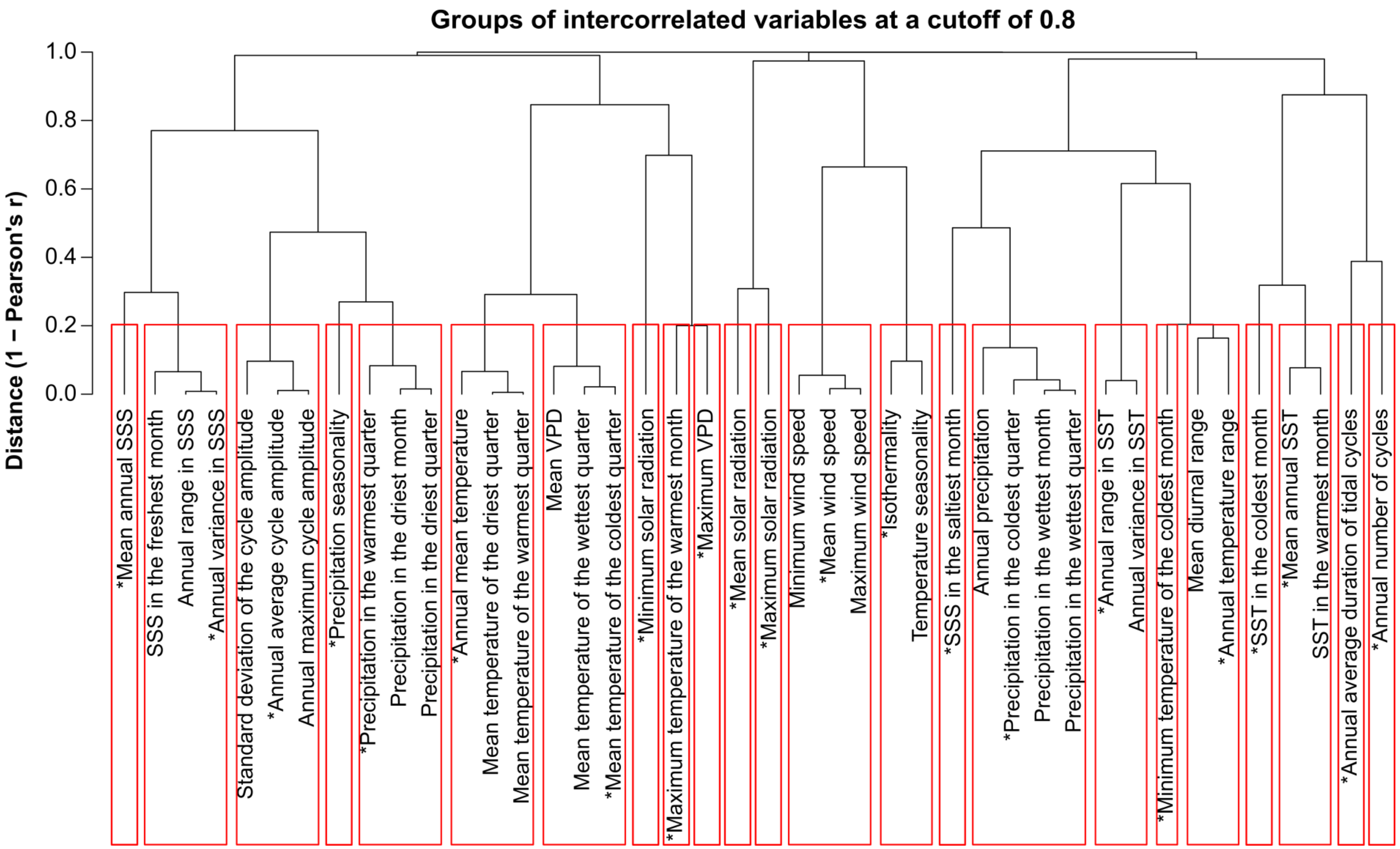


**Figure S1.** Groups (red boxes) of correlated environmental variables retrieved from the public data platforms WorldClim (Fick & Hijmans, 2017), Marspec (Sbrocco & Barber, 2013), and ECDS (Klein *et al.*, 2013). The cutoff value for Pearson's correlation coefficient was set to 0.8. *Environmental variables retained for subsequent analysis. SST = sea surface temperature. SSS = sea surface salinity. VPD = Vapor pressure deficit.

**REFERENCES**


Fick SE, Hijmans RJ. 2017. WorldClim 2: new 1-km spatial resolution climate surfaces for global land areas. *International Journal of Climatology* 37: 4302-4315.

Klein T, Langner J, Frankenberg B, Svensson J, Broman B, Bennet C, Langborg T. 2013. ECDS - a Swedish research infrastructure for the open sharing of environment and climate data. *Data Science Journal* 12: 1-9.

Sbrocco EJ, Barber PH. 2013. MARSPEC: ocean climate layers for marine spatial ecology. *Ecology* 94: 979.
